# Nurseries and garden centres as hubs of alien plant invasions

**DOI:** 10.1101/2024.10.04.616618

**Authors:** Judit Sonkoly, V Attila Molnár, Péter Török, Kristóf Süveges, Attila Takács

## Abstract

The growing global horticultural trade is having a steadily increasing impact on the rate at which alien species are introduced into new areas, partly because horticultural trade also entails the unintentional dispersal of many contaminant species. Although there are reports about noteworthy occurrences of alien plant species in garden centres, this phenomenon has hardly been studied systematically. To bridge this knowledge gap, we conducted systematic field surveys in 12 garden centres in Hungary to assess their alien flora. We hypothesised that (i) the number of alien species inhabiting a garden centre is positively correlated with its size, (ii) relative to their size, garden centres host a disproportionately large proportion of the local alien flora, and (iii) alien species inhabiting garden centres differ from the regional alien flora in their traits. We recorded altogether 93,788 individuals of 67 introduced species, seven of which have not yet been reported from the country. There was considerable variability in the number of species and individuals found in each garden centre, but there was no correlation between the size of the garden centres and the number of species they host. Despite their relatively small size, the studied garden centres hosted a considerable proportion of the local alien flora, indicating that they strongly accumulate alien species and that they can act as invasion hubs for several alien species. Alien species inhabiting garden centres differed from the regional alien flora in some of their trait values, indicating that the species that are most successful at establishing populations inside garden centres are both good dispersers and possess an effective resource-acquisitive strategy. We conclude that established alien plant populations in garden centres may induce local invasions, and in the meantime, individuals and seeds inside the containers of ornamental plants are regularly transported to distant areas by the customers. Therefore, plant species dispersed as contaminants of horticultural stock need to be better considered in invasion biology to reduce the threat they may present.

## Introduction

Invasive alien species (IAS) are one of the major drivers of environmental change worldwide (Dí;az et al. 2019). As summarised by the review of Pyšek et al. (2020), IAS can have a multitude of adverse effects, such as reducing biodiversity, disrupting food webs, or altering habitat structure, ultimately altering ecosystem functions and services (Ehrenfeld 2010, Vilà et al. 2011). Through all these profound impacts, IAS can also be detrimental to human health and economy (Early et al. 2016, Haubrock et al. 2021). Global changes including climate change and anthropogenic habitat conversion are the major drivers of the spreading of IAS (Bradley et al. 2010, Pyšek et al. 2020), and climate change is expected to further increase the rate and impact of plant invasions globally (Early et al. 2016, Dullinger et al. 2017). Hence, gaining a thorough understanding of the spread of IAS is ever more important.

Despite being considered rare and stochastic, long-distance dispersal (LDD) events play a crucial role in multiple large-scale ecological processes (Nathan 2006, Paulose et al. 2020). Besides gene flow between distant populations (Clobert et al. 2012), the long-term persistence of species in fragmented habitats (Trakhtenbrot et al. 2005), and range shifts to keep up with changing landscapes and climates (Clark et al. 1999), biological invasions are also mainly driven by LDD events and not by local dispersal (Kot et al. 1996, Nathan 2006). The increasing recognition of the exceptional importance of LDD in several processes of great conservation concern has resulted in growing numbers of studies of this phenomenon. However, as LDD is very difficult to study, our understanding of it is still severely limited (Levin et al. 2003, Jordano 2017), which also prevents us from fully understanding the processes underlying the spread of IAS. Human activity has historically been a considerable factor in the dispersal of species, both accidentally and on purpose (Hulme 2009, Bullock et al. 2018). Moreover, due to the quickly rising rates of global trade and human mobility in the last century or so, humanity has presumably become the major dispersal vector (Hodkinson & Thompson 1997). Humanity can now also be considered as the main LDD vector (Higgins et al. 2003, Taylor et al. 2012), suggesting that the increasing levels of globalisation and human activity has led to more chances for LDD events to occur.

Likely associated with the increased chances of LDD events, all evidence suggests that the pace of alien species accumulation in various locations is also increasing due to the progression of free trade and globalization (Lockwood et al. 2005, Hulme 2009), and the spread of IAS is widely considered to be mainly driven by international trade and transport both in terrestrial and in aquatic ecosystems (Jenkins 1996, Perrings et al. 2005, Essl et al. 2015). Studies have demonstrated a clear link between the monetary value of imports and the number of alien species in the country (Vilà & Pujadas 2001, Westphal et al. 2008, Dalmazzone & Giaccaria 2014), and the rate of alien species accumulation of different species groups in the United States was well described by a model using the cumulative value of imports (Levine & d’Antonio 2003).

Horticulture has always been one of the most significant human activities introducing alien species into new areas (Kenis et al. 2007, van Kleunen et al. 2018), but the growing global horticultural trade (van Kleunen et al. 2018) is having a steadily increasing impact on the rate at which alien species are introduced into new areas (Bradley et al. 2012, Liebhold et al. 2012). Planted ornamental species are widely known as the main source of introduced alien plants, and many of them are capable of becoming naturalised and subsequently invasive in their introduced range (Haeuser et al. 2018, Hulme et al. 2018). However, the global horticultural trade also entails the unintentional dispersal of many contaminant species (Hodkinson & Thompson 1997, Montagnani et al. 2022, Sonkoly et al. 2022).

Nurseries and garden centres (hereafter garden centres for simplicity) have been repeatedly shown to accumulate a wide range of introduced species, such as plants (e.g., Hoste et al. 2009, Gallego and Lumbreras 2013, Takács et al. 2020, Rigó et al. 2023), plant pathogens (Bienapfl & Balci 2014, Jung et al. 2016), molluscs (Bergey et al. 2014, Páll-Gergely et al. 2020), or planarians (Lazányi et al. 2024). Alien species found in the diverse anthropogenic microhabitats inside the garden centres can arrive there through various pathways. They can easily infiltrate from existing populations nearby the establishments, but they may arrive from much more distant locations via LDD, for example, they may be dispersed on the clothing and footwear of the staff or the costumers (Ansong & Pickering 2016, or Lukács et al. 2024) or attached to vehicles (Taylor et al. 2012, von der Lippe & Kowarik 2007). Moreover, as horticultural substrates are frequently contaminated with seeds and other plant propagules (Conn et al. 2008, Dyer et al. 2017, Sonkoly et al. 2022), imported container plants can be expected to disperse species over large distances and potentially introduce new alien species into garden centres.

To develop evidence-based and effective strategies for the prevention of further plant invasions, more research attention needs to be directed to the role of horticultural trade in the spread of IAS (van Kleunen et al. 2018). Although there are reports about noteworthy occurrences of alien plant species in garden centres (e.g., Gallego and Lumbreras 2013, Takács et al. 2020, Rigó et al. 2023), this phenomenon has hardly been studied systematically (but see Hoste et al. 2009). To bridge this knowledge gap, we conducted systematic field surveys assessing which alien plant species occur in garden centres and in what abundance. We hypothesised that (i) the number of alien species inhabiting a garden centre is positively correlated with the size of the given garden centre, (ii) relative to their size, garden centres host a disproportionately large proportion of the local alien flora, and (iii) alien species inhabiting garden centres differ from the regional alien flora in their traits related to dispersal and ecological strategy.

## Materials & Methods

In 2019, we conducted three surveys (spring, summer, and autumn, separated by 2-3-month intervals) at 12 ornamental plant retailers in and near the city of Debrecen, Hungary (see Fig. 1). The surveyed retailers included both nurseries and garden centres of different sizes from small, family-owned businesses to large franchise retailers (ranging from cc. 1200 m^2^ to cc. 12,000 m^2^ in area). Owners and/or managers were informed and gave their consent to conduct the surveys.

**Figure 1.**
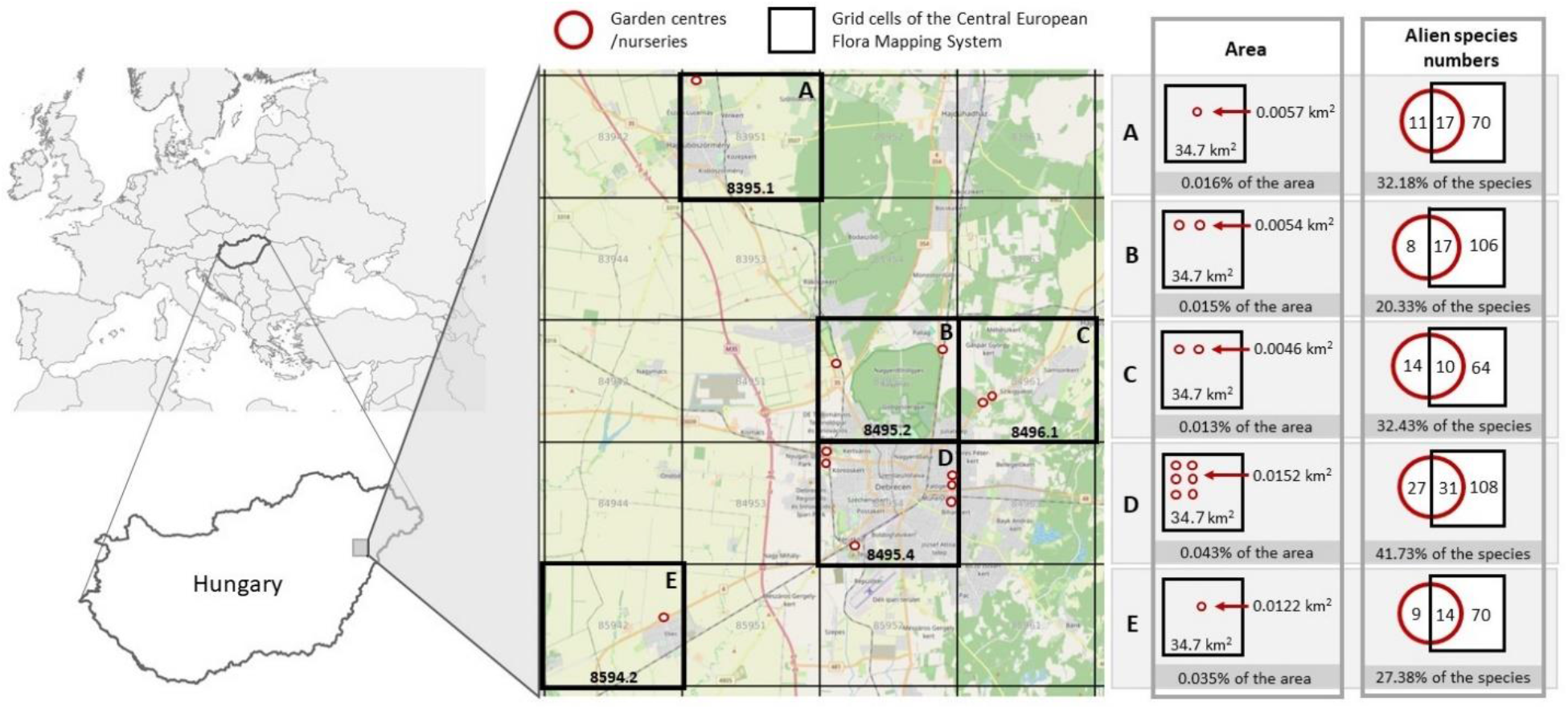
The location of the studied garden centres in and around the city of Debrecen, Hungary. The panel on the right shows the area of the garden centres in each quarter-grid cell of the Central European Flora Mapping System (Niklfeld 1971) compared to the total area of the grid cell, together with the number of alien plant species we found in the garden centres, the documented number of alien plant species in the respective grid cell according to the Vascular plants of Hungary online database (Bartha et al. 2021+), and the overlap between the two sets of species.

Repeated surveys in each season during the vegetation period ensured the finding of species with different phenology. We surveyed the whole territory of each garden centre including both outdoor and indoor areas (buildings and greenhouses), examining the pots and containers of ornamentals for sale, as well as flowerbeds, pioneer or covered soil surfaces (with agrotextile or gravel), pavement cracks and fugues, lawns and other vegetated areas. We recorded all spontaneously occurring individuals of all alien plant species, therefore excluding planted individuals and the plants offered for sale. We recorded ornamental plant species (e.g., *Buddleja davidii, Pilea microphylla, Soleirolia soleirolii*) only if individuals of them were established spontaneously (as a contaminant) in pots of other species or outside their pots. For species that spread clonally or form mats and in the case of extremely abundant species the number of individuals was estimated. The native versus alien status of species in Hungary was assessed using the checklist of Csiky et al. (2023); neophytes and archaeophytes were both treated as aliens. We treated *Oxybasis glauca* (whose native or alien status in Hungary is debated) as an alien species and therefore included it in the analyses.

The area of the garden centres was measured using Google Earth. The correlation between the area of each garden centre and the number of species and individuals they host was tested using Pearson correlations. The studied garden centres were scattered in five quarter grid cells (hereafter grid cells for simplicity) of the Central European Flora Mapping System (Niklfeld 1971, see Fig. 1). We compared the alien species list of the garden centres found in each grid cell with the list of the documented alien species in the same grid cell according to Bartha et al. (2021+). To test whether garden centres host a disproportionately large proportion of the local alien flora relative to their size, we calculated the percentage of each grid cell’s area covered by the garden centre(s) within it, and also the percentage of the alien species of each grid cell (also including the new species found by our survey) found within the area of the garden centre(s) within the given grid cell. The percentages were compared by Wilcoxon rank sum test.

When comparing traits and ecological indicator values (EIVs) of the species found in the garden centres with the traits and EIVs of the total regional alien flora, we considered the alien checklist of Csiky et al. (2023) and obtained data for them from the Pannonian Database of Plant Traits (PADAPT, Sonkoly et al. 2023). We considered those traits which may influence the species’ ability to establish in the garden centres and for which data was available for the majority of the species. Based on these criteria, we analysed thousand-seed mass (TSM, g), plant height (cm), leaf area (LA, mm^2^), specific leaf area (SLA, mm^2^/mg), leaf dry matter content (LDMC, mg/g), and leaf dry weight (LDW, mg). As EIVs, we considered the Borhidi-type indicator values (Borhidi 1995) for soil moisture (W_B_), light intensity (L_B_) and nutrient supply (N_B_), which are based on the indicator values of Ellenberg (Ellenberg et al. 1992) and also adapted to the Pannonian Ecoregion. In case of the species documented in the garden centres, traits and ecological indicator values were also primarily obtained from the regional database PADAPT (Sonkoly et al. 2023), but missing trait values were filled in wherever possible, either by new measurements carried out for the purpose of this analysis or by gathering data from the LEDA Traitbase (Kleyer et al. 2008). Traits and EIVs of the two species groups (i.e. recorded aliens vs. the regional alien flora) were compared using Welch’s t-tests. All statistical analyses were conducted in an R environment (version 4.3.2, R Core Team 2023). Nomenclature follows the Euro+Med PlantBase (2006+).

## Results

We recorded altogether 93,788 individuals of 67 introduced species (see Supplementary Material 1), seven of which have not yet been reported from Hungary: *Acalypha brachystachya, Fumaria capreolata, Gnaphalium pensylvanicum, Hylotelephium sieboldii, Oldenlandia corymbosa, Soleirolia soleirolii*, and *Urtica pilulifera*. The five most abundant species were *Sagina procumbens, Oxalis corniculata, Cardamine occulta, Oxybasis glauca*, and *Euphorbia maculata*, each with more than 10 thousand documented individuals. On the other hand, 13 species (19%) were each represented by a single individual (Supplementary Material 1). On average, each species occurred in four of the 12 studied garden centres. Twenty-eight species (42%) were found in a single one, while six species (*Erigeron annuus, Erigeron canadensis, Euphorbia maculata, Oxalis corniculata, Oxalis dillenii*, and *Sagina procumbens*) occurred in all of the garden centres (Supplementary Material 1). Fifty-two of the 67 documented species occurred in the containers of the ornamental plants. There were 15 species occurring only outside containers, while 15 species occurred only inside the containers.

The studied garden centres each hosted an average of 7,816 individuals of 22 alien species. However, considerable variability was observed between the garden centres, with species numbers ranging from 11 and 33 and the total number of individuals ranging between 1,440 and 24,877. The observed number of species in a garden centre was not correlated with the area of the given garden centre (*p*=0.901), however, the number of individuals was significantly positively correlated with it (*R*=0.817, *p*=0.001). The number of species and the number of individuals in a garden centre were also not correlated (*p*=0.808).

The studied garden centres were scattered in five grid cells of the Central European Flora Mapping System (see Fig. 1). We found that although garden centres covered a very small fraction of the grid cells (ranging from 0.013% to 0.043%), they encompassed surprisingly high alien species numbers. On average, 17.2% (ranging from 13.5% to 22.3%) of the already known alien flora of the grid cells was found in the comparatively tiny area of the garden centres. In addition, we found 8 to 27 species in the garden centres of each grid cell that had not been previously documented in those grid cells (Fig. 1). The percentage of the alien species of each grid cell found within the area of the garden centre(s) within the given grid cell was substantially higher than the percentage of each grid cell’s area covered by the garden centre(s) within it (W=0, p=0.007, Wilcoxon rank sum test), demonstrating that garden centres host a disproportionately large proportion of the local alien flora relative to their size (Fig. 1).

When we compared the alien species documented in the garden centres with the regional alien flora in terms of some traits and ecological indicator values (EIVs) we found that the alien species documented in the garden centres had significantly lighter seeds, higher specific leaf area (SLA), lower leaf dry matter content (LDMC), and higher EIVs for soil moisture and nutrient supply than the regional alien flora in general (Table 1).

**Table 1.** Traits and ecological indicator values (EIVs) of the alien species documented in the garden centres compared to the traits and EIVs of the regional alien flora (Welch’s t-tests). Abbreviations: TSM – thousand-seed mass; LA – leaf area; SLA – specific leaf area; LDMC – leaf dry matter content; LDW – leaf dry mass; W_B_, L_B_, and N_B_ – Ecological indicator values for soil moisture, light intensity and nutrient supply, respectively (Borhidi 1995).

|  | Regional alien flora<br>(mean±SE) | Garden centres<br>(mean±SE) | *p* value |
| --- | --- | --- | --- |
| **TSM (g)** | 30.07±6.10 | 10.51±4.26 | **0.009** |
| **Plant height (cm)** | 306.14±24.19 | 434.66±114.07 | 0.275 |
| **LA (mm<sup>2</sup>)** | 3132.30±385.55 | 2479.47±675.4 | 0.403 |
| **SLA (mm<sup>2</sup>/mg)** | 23.50±0.75 | 31.04±2.33 | **0.003** |
| **LDMC (mg/g)** | 247.54±5.21 | 212.90±11.67 | **0.010** |
| **LDM (mg)** | 196.41±28.49 | 119.84±33.10 | 0.081 |
| **$W_B$** | 4.54±0.08 | 5.11±0.21 | **0.015** |
| **$L_B$** | 7.39±0.05 | 7.17±0.19 | 0.263 |
| **$N_B$** | 5.37±0.10 | 6.53±0.25 | **<0.0001** |

## Discussion

The role of horticultural trade in biological invasions has been mainly considered in the context of ornamental plants escaping cultivation and becoming invasive, and it has been much less considered as a pathway of accidental human-vectored dispersal (Bullock et al. 2018) of contaminant species. Here, we demonstrated that, despite their small area, garden centres accumulate alien plant species in surprisingly large numbers, and that a considerable proportion of these species had large established populations which can be starting points of local dispersal in the surrounding landscape. The fact that the majority of the species were found inside the containers of the ornamental plants supports our assumption that most accidental alien plant species arrive at the garden centres via shipments of ornamental plants, thus by long-distance dispersal (LDD). This is also supported by the presence of species that have not been previously documented in Hungary.

The seven species documented in Hungary for the first time fall into three categories: (i) Species of Mediterranean origin, which presumably arrived in the garden centres due to the increasing involvement of Mediterranean countries, particularly Italy and Spain, in the container plant trade within Europe (see Hoste & Verloove 2009): *Fumaria capreolata* and *Urtica pilulifera*. (ii) Emerging species of (sub)tropical origin, which are not widespread yet, but have already been introduced to some temperate regions: *Acalypha brachystachya, Gnaphalium pensylvanicum*, and *Oldenlandia corymbosa*. (iii) Ornamental species, which are probably spreading in the garden centres due to their widespread cultivation: *Hylotelephium sieboldii* and *Soleirolia soleirolii*. Seventeen of the 67 species we documented were also found in containers of ornamental plants in Belgium (Hoste et al. 2009), including two of the species new to the Hungarian flora (*F. capreolata* and *G. pensylvanicum*).

The most abundant alien species in the garden centres were mostly already quite widespread both in Hungary in Europe in general (Kalusová et al. 2024). However, the third most abundant species, *Cardamine occulta*, of which we documented more than 11,000 individuals, had only been included in the alien plant inventory of eight out of 55 European countries and territories up until 2022 (Kalusová et al. 2024), and it was also only recently found in Hungary (Takács et al. 2020). This species seems to quickly spread in Europe in places where its occurrence can be attributed to the horticultural trade (see e.g., Leostrin & Mayorov 2019, Šlenker et al. 2019, Pliszko 2020, Hruševar et al. 2021, Király et al. 2021, Kovács et al. 2023).

Both the number of species and the number of individuals were highly variable across the studied garden centres. We hypothesised that the number of alien species found in a garden centre is positively related to the size of the given garden centre, as larger garden centres presumably have a larger incoming stock. A larger incoming stock can be associated with higher propagule pressure (i.e., a larger number of individuals, for example propagules, released into the area), which is considered as one of the most important factors determining the success of invasive species (Lockwood et al. 2005, Colautti et al. 2006). Moreover, surveying land snails in plant nurseries, Bergey et al. (2014) found that the number of taxa occurring in a nursery was positively related to the nursery area. Despite this, there was no relationship between the size of a garden centre and the number of alien plant species it hosts. However, there was a significant positive relationship between the size of a garden centre and the number of alien plant individuals found in it. The highest numbers of alien species were observed in the garden centres that, rather than being the largest in area, were the busiest and offered the greatest diversity of plants for sale based on our experience. This observation is in line with the key importance of propagule pressure, but, unfortunately, we have no data to test this assumption.

We also hypothesised that, relative to their size, garden centres host a disproportionately large proportion of the local alien flora. We demonstrated that although the studied garden centres cover only a very small fraction of the area of the flora mapping grid cells they are located in, they host a considerable proportion of the documented alien flora within those grid cells, confirming our hypothesis. This result illustrates that garden centres strongly accumulate alien species and that they can act as invasion hubs for several new invading species (Letnic et al. 2015). Human-vectored dispersal can result in spatially aggregated populations of alien species, which are regularly focused in and around anthropogenic habitats (Bullock et al. 2018). These habitats can subsequently function as invasion hubs, from which established satellite populations of alien species can spread further into the landscape (Suarez et al. 2001, With 2002). Because eradicating such invasion hubs (or ‘nascent foci’) has been shown to be a targeted and effective approach to hinder the spread of IAS (Moody & Mack 1988, Florance et al. 2011), identifying invasion hubs is a very important step towards controlling the spread of IAS.

Our hypothesis that alien species inhabiting garden centres differ from the regional alien flora in their traits was partially supported by the results. We found no difference between the two species groups in terms of plant height and leaf size (expressed in terms of specific leaf area and leaf dry mass). On the other hand, the documented alien species had significantly lower LDMC and higher SLA compared to the regional alien flora, which indicates that they are placed on the leaf economic spectrum towards the more fast-growing, resource-acquisitive strategies (Leishman et al. 2007). The fact that the documented alien species also had lower seed mass is probably due to the presumed better dispersal ability (Guo et al. 2000, Tremlová & Münzbergová 2007) and longer seed persistence (Thompson et al. 1993, Zhao et al. 2011) of small-seeded species. Having persistent seeds ensures that seeds can remain viable in the containers throughout the potentially lengthy transportation process from wholesalers to garden centres (and eventually to the costumers’ gardens). These findings indicate that the species that are most successful at establishing populations inside garden centres are both good dispersers and possess a resource-acquisitive strategy (Wright et al. 2004, Pierce et al. 2014), because they need to be able to grow and reproduce quickly. Species with slow growth rate, high stature or long life cycle also occasionally appear in garden centres, but they are less abundant and unlikely to establish self-sustaining populations within the garden centres. Thus, horticultural trade can play a role in their introduction, but the garden centres themselves play a less important role as invasion hubs for these species (Suarez et al. 2001, Letnic et al. 2015). Regrading ecological indicator values, we found that the alien species inhabiting the garden centres had significantly higher values for soil moisture and nutrient supply compared to the regional alien flora, indicating the effect of fertilizer use and frequent irrigation in nurseries and garden centres.

Fifty-two of the identified 67 species occurred in containers of ornamental plants. As the customers regularly transport the containers over great distances, further LDD of these species can be expected regularly (Hodkinson & Thompson 1997, Bergey et al. 2014). The 15 species found only outside the containers may have dispersed into the garden centres from existing populations in the neighbouring area, by other means of dispersal rather than as contaminants of horticultural stock. Individuals outside containers are not readily transported to distant areas, but as several species had established populations of considerable size even outside the containers (*Cardamine occulta, Euphorbia* spp., *Oxalis* spp., *Veronica peregrina* etc.), they can exert sufficient propagule pressure to induce local invasions.

Regarding the genetic effects of accidental LDD by the horticultural trade, it has to be considered that LDD events commonly lead to low genetic diversity of the introduced populations due to strong founder effects (Setsuko et al. 2020, Wu et al. 2023). On the other hand, LDD by humans often transports many propagules or individuals of a species at a time (Bullock et al. 2018). If propagules arriving at a site are from different source areas and the genetic makeup of the species in its native range is spatially structured, the genetic diversity of such introduced populations may be higher than that of any of the native source populations (Lockwood et al. 2005, Smith et al. 2020), which can minimize Allee effects and founder effects (Wu et al. 2023). These considerations are in line with the notion that the international horticultural trade not only has a key role in new species introductions, it also considerably increases the chance of successful invasions due to the recurring introductions of alien plants into local outlets of the global horticultural trade (Dehnen-Schmutz et al. 2007), which increase the chances of novel genetic combinations and evolutionary shifts (Kolbe et al. 2004, Lockwood et al. 2005, Šlenker et al. 2018).

It has been repeatedly suggested that managing and controlling invasive species is the easiest and most cost-effective in the early stages of the invasion process (e.g., Rout et al. 2011, Simberloff et al. 2013, Chapman et al. 2016). Specifically, eradicating recently established satellite populations (Moody and Mack 1988, Florance et al. 2011) or reducing the probability of LDD events that create such populations (Buckley et al. 2005, Shea et al. 2010) can be especially effective approaches to control the spread of IAS. The above considerations suggest that eradicating the newly established populations of alien species in garden centres would be easier and more cost-effective in controlling plant invasions than focusing the control measures on older, and thus bigger populations. Therefore, early detection and eradication of these satellite populations in garden centres should be a priority in IAS management. In the meantime, we need to raise public awareness, thus both the managers and employees of plant nurseries and garden centres, as well as the customers should be educated and informed about the potential adverse consequences of new plant species introduced as contaminants of container plants.

It has to be noted that our assessment of the role of garden centres as invasion hubs is somewhat limited due to the lack of reliable information about which species actually arrived at the garden centres by LDD via the horticultural trade and which species dispersed into them from existing populations in the neighbouring areas. However, it does not refute the fact that even the species originally infiltrating the garden centres from the neighbouring populations can be further dispersed by the customers transporting the containers to distant areas. Due to these limitations and the complexity of this phenomenon, many questions about the issue remain unanswered. For example, the propagule content of the containers of ornamental plants has been largely understudied so far (but see Cross & Skroch 1992, and Conn et al. 2008). Assessing the different pathways by which propagules can reach these containers could also provide information about the step of the horticultural trade at which interventions could be the most effective.

## Conclusions

Our study is one of the few assessments of the accidental dispersal of plants as contaminants of the horticultural trade. We showed that garden centres host a large number of accidental alien species, and as many species had large established populations within the garden centres, they can serve as invasion hubs for new IAS. Established populations within the garden centres can exert sufficient propagule pressure to induce local invasions. In the meantime, individuals established in containers and their seed bank inside them are regularly transported to distant areas by the customers, where they may form new invasion hubs. Thus, horticultural trade is not only a pathway of alien plant introductions, it can also play a role in the subsequent stages of the invasion process. Due to the diverse range of species and the huge amount of propagules possibly arriving accompanying the increasing inflow of horticultural stock, it is highly likely that at least some of them will be naturalised and ultimately become invasive in their new range. Although more attention has been paid to other introduction pathways such as deliberate introductions for ornamental purposes, our study demonstrated that plant species dispersed as contaminants of horticultural stock need to be better considered in invasion biology to reduce the threat they may present.

## Acknowledgements

We are grateful for Edina Tóth-Szabó and Balázs Hubicska for their help in carrying out the surveys, and for Andrea McIntosh-Buday for her help with some trait measurements. The authors are thankful for the support of the National Research, Development and Innovation Office (PD 137747, KKP144068; K137573; K132573) and to the Bolyai János Scholarship of the Hungarian Academy of Sciences (BO/00587/23/8) during manuscript preparation. Supported by the ÚNKP-23-5 New National Excellence Program of the Ministry for Culture and Innovation from the source of the National Research, Development and Innovation Fund.

