## Supplementary Information (Table S1 and S2) for "Nurseries and garden centres as hubs of alien plant invasions"

**Table S1.** The number of individuals documented in the studied garden centers for each species, and their invasion status in Europe and Hungary.

| Species | Total number of individuals | Individuals inside containers | Individuals outside containers | Number of garden centers | Invasion status in Europe (Kalusová et al. 2024) | Invasion status in Hungary (Csiky et al. 2023) |
|---|---|---|---|---|---|---|
| *Acalypha brachystachya* | 17 | 15 | 2 | 2 | *not listed yet* | *not listed yet* |
| *Acer negundo* | 1 | 1 | 0 | 1 | invasive | naturalized |
| *Acer saccharinum* | 200 | 200 | 0 | 1 | naturalized | naturalized |
| *Ailanthus altissima* | 4 | 1 | 3 | 4 | invasive | transformer |
| *Amaranthus blitum* | 94 | 51 | 43 | 10 | invasive | naturalized |
| *Amaranthus deflexus* | 37 | 9 | 28 | 7 | invasive | naturalized |
| *Amaranthus powellii* | 186 | 2 | 184 | 3 | invasive | naturalized |
| *Amaranthus retroflexus* | 412 | 26 | 386 | 7 | invasive | naturalized |
| *Ambrosia artemisiifolia* | 827 | 91 | 736 | 8 | invasive | naturalized |
| *Artemisia annua* | 1 | 0 | 1 | 1 | invasive | invasive |
| *Asclepias syriaca* | 1 | 0 | 1 | 1 | invasive | transformer |
| *Buddleja davidii* | 10 | 0 | 10 | 1 | invasive | naturalized |
| *Cardamine occulta* | 11,395 | 7,642 | 3,753 | 11 | naturalized | *not listed yet* |
| *Chenopodiastrum hybridum* | 3 | 0 | 3 | 2 | naturalized | naturalized |
| *Chenopodiastrum murale* | 8 | 5 | 3 | 1 | naturalized | naturalized |
| *Chenopodium album* | 95 | 21 | 74 | 7 | naturalized | naturalized |
| *Chenopodium betaceum* | 33 | 13 | 20 | 6 | casual | naturalized |
| *Chenopodium ficifolium* | 165 | 164 | 1 | 4 | naturalized | naturalized |

| | | | | | | |
|---|---|---|---|---|---|---|
| *Claytonia perfoliata* | 1 | 1 | 0 | 1 | naturalized | casual |
| *Commelina communis* | 4 | 0 | 4 | 1 | invasive | naturalized |
| *Dysphania pumilio* | 1 | 1 | 0 | 1 | invasive | naturalized |
| *Eclipta prostrata* | 8 | 3 | 5 | 4 | invasive | *not listed yet* |
| *Elaeagnus angustifolia* | 1 | 1 | 0 | 1 | invasive | transformer |
| *Epilobium ciliatum* | 228 | 147 | 81 | 7 | invasive | naturalized |
| *Erigeron annuus* | 4,529 | 1,498 | 3,031 | 12 | invasive | invasive |
| *Erigeron canadensis* | 1,629 | 528 | 1,101 | 12 | invasive | invasive |
| *Erigeron sumatrensis* | 52 | 0 | 52 | 2 | invasive | *not listed yet* |
| *Euphorbia maculata* | 10,113 | 4,352 | 5,761 | 12 | invasive | invasive |
| *Euphorbia peplus* | 758 | 109 | 649 | 8 | invasive | naturalized |
| *Euphorbia prostrata* | 167 | 11 | 156 | 4 | invasive | naturalized |
| *Euphorbia serpens* | 3550 | 918 | 2,632 | 5 | invasive | naturalized |
| *Fumaria capreolata* | 1 | 1 | 0 | 1 | naturalized | *not listed yet* |
| *Galinsoga parviflora* | 726 | 28 | 698 | 8 | invasive | invasive |
| *Galinsoga quadriradiata* | 73 | 47 | 26 | 5 | invasive | naturalized |
| *Geranium dissectum* | 2 | 0 | 2 | 1 | naturalized | naturalized |
| *Gleditsia triacanthos* | 105 | 0 | 105 | 1 | invasive | naturalized |
| *Gnaphalium pensylvanicum* | 1 | 1 | 0 | 1 | invasive | *not listed yet* |
| *Humulus japonicus* | 1 | 0 | 1 | 1 | invasive | invasive |
| *Hylotelephium sieboldii* | 1 | 1 | 0 | 1 | naturalized | *not listed yet* |
| *Lepidium didymum* | 2 | 2 | 0 | 1 | invasive | casual |
| *Microrrhinum minus* | 21 | 0 | 21 | 1 | invasive | naturalized |
| *Oenothera biennis* | 3 | 0 | 3 | 2 | invasive | invasive |
| *Oldenlandia corymbosa* | 26 | 26 | 0 | 2 | *not listed yet* | *not listed yet* |
| *Oxalis corniculata* | 15,662 | 8,614 | 7,048 | 12 | invasive | invasive |

| | | | | | | |
|---|---|---|---|---|---|---|
| *Oxalis dillenii* | 6,011 | 1,789 | 4,222 | 12 | invasive | invasive |
| *Oxalis stricta* | 3,640 | 2,239 | 1,401 | 4 | invasive | invasive |
| *Oxybasis glauca* | 11,197 | 4,314 | 6,883 | 9 | naturalized | naturalized |
| *Panicum dichotomiflorum* | 15 | 0 | 15 | 1 | invasive | invasive |
| *Panicum riparium* | 44 | 36 | 4 | 2 | casual | casual |
| *Parietaria judaica* | 267 | 8 | 259 | 3 | naturalized | naturalized |
| *Parthenocissus quinquefolia* | 3 | 0 | 3 | 1 | invasive | naturalized |
| *Phytolacca americana* | 1 | 0 | 1 | 1 | invasive | invasive |
| *Pilea microphylla* | 106 | 79 | 27 | 5 | casual | *not listed yet* |
| *Polycarpon tetraphyllum* | 1 | 1 | 0 | 1 | naturalized | naturalized |
| *Polypogon viridis* | 11 | 6 | 5 | 1 | naturalized | casual |
| *Populus × canadensis* | 2 | 2 | 0 | 1 | invasive | naturalized |
| *Potentilla indica* | 2 | 2 | 0 | 1 | invasive | invasive |
| *Robinia pseudoacacia* | 12 | 11 | 1 | 2 | invasive | transformer |
| *Sagina procumbens* | 17,672 | 4,714 | 12,958 | 12 | naturalized | naturalized |
| *Senecio leucanthemifolius ssp. vernalis* | 6 | 0 | 6 | 1 | invasive | naturalized |
| *Soleirolia soleirolii* | 5 | 5 | 0 | 1 | invasive | *not listed yet* |
| *Solidago canadensis* | 36 | 9 | 27 | 3 | invasive | transformer |
| *Solidago gigantea* | 207 | 139 | 68 | 4 | invasive | transformer |
| *Ulmus pumila* | 196 | 78 | 118 | 5 | invasive | invasive |
| *Urtica pilulifera* | 1 | 1 | 0 | 1 | naturalized | *not listed yet* |
| *Veronica peregrina* | 2,787 | 626 | 2,161 | 9 | invasive | naturalized |
| *Veronica persica* | 412 | 4 | 408 | 6 | invasive | naturalized |

**Table S2.** Traits and ecological indicator values of the species documented in the studied garden centers. Abbreviations: TSM – thousand-seed mass; LA – leaf area; SLA – specific leaf area; LDMC – leaf dry matter content; LDM – leaf dry mass; $W_B$ – Borhidi-type indicator value for soil moisture (Borhidi 1995); $L_B$ – Borhidi-type indicator value for light intensity (Borhidi 1995); $N_B$ – Borhidi-type indicator value for nutrient supply (Borhidi 1995). Trait data were gathered primarily from a regional database (PADAPT, Sonkoly et al. 2023), but missing trait values were filled in wherever possible, either by new measurements carried out for the purpose of this analysis or by gathering data from the LEDA Traitbase (Kleyer et al. 2008).

| **Species** | **TSM (g)** | **Plant height (cm)** | **LA ($mm^2$)** | **SLA ($mm^2/mg$)** | **LDMC (mg/g)** | **LDM (mg)** | **$W_B$** | **$L_B$** | **$N_B$** |
|---|---|---|---|---|---|---|---|---|---|
| *Acalypha brachystachya* | NA | NA | NA | NA | NA | NA | NA | NA | NA |
| *Acer negundo* | 41.17 | 2000 | 7417.88 | 20.20 | 296.72 | 357.17 | 6 | 5 | 7 |
| *Acer saccharinum* | 126.06 | 3000 | 8233.10 | 14.09 | 368.30 | 584.50 | NA | NA | NA |
| *Ailanthus altissima* | 33.38 | 1500 | 33971.20 | 17.48 | 234.42 | 1437.67 | 5 | 6 | 8 |
| *Amaranthus blitum* | 0.34 | 80 | 427.15 | 26.84 | 215.37 | 15.00 | 4 | 8 | 8 |
| *Amaranthus deflexus* | 0.31 | 50 | 747.80 | 29.56 | 155.21 | 25.30 | 4 | 8 | 7 |
| *Amaranthus powellii* | 0.49 | 100 | 1283.97 | 17.17 | 214.85 | 74.77 | 4 | 9 | 7 |
| *Amaranthus retroflexus* | 0.398 | 100 | 1486.97 | 28.82 | 183.67 | 55.97 | 5 | 9 | 9 |
| *Ambrosia artemisiifolia* | 5.66 | 150 | 588.46 | 29.73 | 257.07 | 34.37 | 5 | 9 | 7 |
| *Artemisia annua* | 0.16 | 150 | 648.94 | 38.82 | 214.80 | 17.33 | 4 | 8 | 6 |
| *Asclepias syriaca* | 5.86 | 150 | 5407.56 | 17.17 | 190.82 | 364.75 | 4 | 7 | 4 |
| *Buddleja davidii* | 0.06 | 500 | 4071.70 | 10.19 | 373.11 | 399.60 | NA | NA | NA |
| *Cardamine occulta* | NA | NA | 262.13 | 38.16 | 132.12 | 6.87 | NA | NA | NA |
| *Chenopodiastrum hybridum* | 1.39 | 60 | 3188.30 | 31.87 | 165.07 | 100.03 | 4 | 7 | 7 |
| *Chenopodiastrum murale* | 0.60 | 60 | 549.30 | 41.39 | 103.67 | 13.27 | 6 | 7 | 7 |
| *Chenopodium album* | 0.76 | 150 | 389.12 | 20.13 | 198.53 | 33.67 | 6 | 8 | 9 |
| *Chenopodium betaceum* | 0.54 | 100 | 558.60 | 16.45 | 235.06 | 33.97 | 6 | 7 | 8 |
| *Chenopodium ficifolium* | 0.26 | 80 | 599.17 | 19.02 | 197.37 | 31.50 | 5 | 8 | 9 |
| *Claytonia perfoliata* | 0.33 | 20 | 964.00 | 79.71 | 36.65 | 12.17 | NA | NA | NA |

| | | | | | | | | | |
|---|---|---|---|---|---|---|---|---|---|
| *Commelina communis* | 8.61 | 70 | 768.97 | 41.93 | 118.76 | 18.20 | 6 | 7 | 6 |
| *Dysphania pumilio* | 0.06 | 50 | NA | NA | NA | NA | NA | NA | NA |
| *Eclipta prostrata* | NA | NA | 1238.20 | 24.23 | 181.21 | 51.10 | NA | NA | NA |
| *Elaeagnus angustifolia* | 122.51 | 1000 | 559.90 | 15.29 | 384.41 | 36.27 | 4 | 8 | 4 |
| *Epilobium ciliatum* | NA | 80 | 659.80 | 56.39 | 112.50 | 11.7 | NA | NA | NA |
| *Erigeron annuus* | 0.04 | 120 | 316.48 | 32.60 | 201.85 | 10.13 | NA | NA | NA |
| *Erigeron canadensis* | 0.06 | 100 | 169.23 | 30.40 | 222.87 | 5.77 | NA | NA | NA |
| *Erigeron sumatrensis* | NA | NA | 430.80 | 22.67 | 171.17 | 19.00 | NA | NA | NA |
| *Euphorbia maculata* | 0.16 | 3 | 17.97 | 15.29 | 380.89 | 1.21 | 5 | 7 | 8 |
| *Euphorbia peplus* | 0.53 | 25 | 125.90 | 91.90 | 137.00 | 1.37 | 7 | 5 | 6 |
| *Euphorbia prostrata* | 0.14 | NA | 56.52 | 27.91 | 226.89 | 2.15 | 3 | 9 | 5 |
| *Euphorbia serpens* | 0.23 | NA | 32.57 | 25.05 | 232.14 | 1.30 | 4 | 6 | 8 |
| *Fumaria capreolata* | 3.67 | NA | NA | NA | NA | NA | NA | NA | NA |
| *Galinsoga parviflora* | 0.14 | 80 | 967.03 | 53.15 | 148.97 | 19.70 | NA | NA | NA |
| *Galinsoga quadriradiata* | 0.14 | 80 | 1398.10 | 74.37 | 102.17 | 18.80 | NA | NA | NA |
| *Geranium dissectum* | 0.73 | 60 | 1025.23 | 26.27 | 182.72 | 40.67 | 6 | 7 | 7 |
| *Gleditsia triacanthos* | 145.62 | 2500 | 5844.03 | 13.67 | 409.94 | 427.57 | 6 | 7 | 8 |
| *Gnaphalium pensylvanicum* | NA | NA | NA | NA | NA | NA | 5 | 7 | 5 |
| *Humulus japonicus* | 9.83 | 500 | 8102.67 | 27.75 | 175.20 | 291.97 | NA | NA | NA |
| *Hylotelephium sieboldii* | NA | NA | 132.80 | 30.18 | 61.97 | 4.40 | NA | NA | NA |
| *Lepidium didymum* | 0.51 | 40 | 158.96 | 35.16 | 148.30 | 7.12 | NA | NA | NA |
| *Microrrhinum minus* | 0.04 | 20 | 46.20 | 17.77 | 172.95 | 2.60 | 5 | 8 | 5 |
| *Oenothera biennis* | 0.46 | 200 | 1880.86 | 17.22 | 301.70 | NA | 3 | 9 | 4 |

| | | | | | | | | | |
|---|---|---|---|---|---|---|---|---|---|
| *Oldenlandia corymbosa* | NA | NA | NA | NA | NA | NA | NA | NA | NA |
| *Oxalis corniculata* | 0.24 | 10 | 363.17 | 61.80 | 128.90 | 6.07 | 4 | 7 | 5 |
| *Oxalis dillenii* | 0.16 | 30 | 297.43 | 29.25 | 203.40 | 10.17 | 5 | 7 | 5 |
| *Oxalis stricta* | 0.21 | 40 | 523.20 | 37.37 | 254.54 | 14.00 | 5 | 7 | 6 |
| *Oxybasis glauca* | 0.21 | 40 | 136.82 | 23.11 | 147.67 | 5.50 | NA | NA | NA |
| *Panicum dichotomiflorum* | 0.9 | 100 | 1424.77 | 19.90 | 338.96 | 59.63 | NA | NA | NA |
| *Panicum riparium* | 0.43 | NA | 3845.67 | 18.91 | 318.20 | 203.33 | NA | NA | NA |
| *Parietaria judaica* | 0.06 | 50 | 645.23 | 70.13 | 146.03 | 9.20 | NA | NA | NA |
| *Parthenocissus quinquefolia* | NA | 3000 | 7824.90 | 29.50 | 186.26 | 265.23 | 5 | 5 | 5 |
| *Phytolacca americana* | 11.88 | 250 | 9736.30 | 56.85 | 128.68 | 171.27 | NA | NA | NA |
| *Pilea microphylla* | NA | NA | 9.01 | 45.04 | 100.00 | 0.20 | NA | NA | NA |
| *Polycarpon tetraphyllum* | 0.0033 | 15 | 11.03 | 26.25 | 262.50 | 0.42 | NA | NA | NA |
| *Polypogon viridis* | NA | NA | NA | NA | NA | NA | NA | NA | NA |
| *Populus × canadensis* | NA | 2250 | 3057.2 | 16.49 | 283.05 | 185.40 | NA | NA | NA |
| *Potentilla indica* | 0.26 | 10 | 1149.50 | 22.09 | 244.11 | 50.16 | 4 | 5 | 8 |
| *Robinia pseudoacacia* | 22.603 | 3000 | 14853.8 | 19.57 | 347.52 | 912.40 | 7 | 6 | 5 |
| *Sagina procumbens* | 0.00 | 5 | 8.76 | 38.40 | 200.55 | 0.23 | NA | NA | NA |
| *Senecio leucanthemifolius ssp. vernalis* | 0.177 | 50 | 193.62 | 21.17 | 132.74 | 18.10 | 4 | 7 | 5 |
| *Soleirolia soleirolii* | NA | 9 | 15.00 | 25.25 | 243.00 | 0.45 | NA | NA | NA |
| *Solidago canadensis* | 0.15 | 180 | 1574.10 | 23.70 | 310.42 | 66.43 | 7 | 7 | 6 |
| *Solidago gigantea* | 0.19 | 150 | 2247.69 | 15.30 | 350.60 | 144.43 | 8 | 7 | 8 |
| *Ulmus pumila* | 8.25 | 1000 | 1195.67 | 9.99 | 416.83 | 119.63 | NA | NA | NA |
| *Urtica pilulifera* | NA | 60 | NA | NA | NA | NA | NA | NA | NA |
| *Veronica peregrina* | 0.04 | 30 | 72.23 | 34.89 | 121.76 | 2.07 | 8 | 8 | 6 |
| *Veronica persica* | 0.25 | 15 | 70.40 | 28.16 | 227.27 | 2.50 | 5 | 6 | 7 |
